# Selective Chemical Labeling and Sequencing of 5-Hydroxymethylcytosine in DNA at Single-Base Resolution

**DOI:** 10.1101/2021.04.13.439733

**Authors:** Xiaogang Li, Xinxin Shi, Yin Gong, Wenting Guo, Yuanrui Liu, Chunwei Peng, Yingchun Xu

**Affiliations:** Department of Clinical Laboratory, State Key Laboratory of Complex Severe and Rare Diseases, Peking Union Medical College Hospital, Chinese Academy of Medical Science and Peking Union Medical College, Beijing, China; Medical Research Center, State Key Laboratory of Complex Severe and Rare Diseases, Peking Union Medical College Hospital, Chinese Academy of Medical Science and Peking Union Medical College, Beijing, China; Beijing Key Laboratory for Mechanisms Research and Precision Diagnosis of Invasive Fungal Diseases, Beijing, China; Gastrointestinal Surgery Department of Gaoxin Branch of the First Affiliated Hospital of Anhui Medical University; School Of Electronics Engineering And Computer Science, Peking University; Characteristic Medical Center of the Chinese People’s Armed Police Force

## Abstract

5-Hydroxymethylcytosine (5hmC), the oxidative product of 5-methylcytosine (5mC) catalyzed by ten-eleven translocation (TET) enzymes, plays an important role in many biological processes as an epigenetic mediator. Prior studies have shown that 5hmC can be selectively labeled with chemically-modified glucose moieties and enriched using click chemistry with biotin affinity approaches. Besides, DNA deaminases of the AID/APOBEC family can discriminate modified 5hmC bases from cytosine (C)or 5-methylcytosine (5mC). Herein, we developed a method based on ESC whole-genome analysis which could enrich 5hmC-containing DNA by selective chemical labeling and locate 5hmC sites at single-base resolution with enzyme-based deamination. The combination experimental design is an extension of previous methods, and we hope that this cost-effective single base resolution 5hmC sequencing method could be used to promote the mechanism and diagnosis research of 5hmC.

## Introduction

5-Hydroxymethylcytosine (5hmC), which exists in various mammalian tissues and cell types, is an oxidative product of 5-methylcytosine (5mC) catalyzed by ten-eleven translocation (TET) enzymes.^1,2^ Many researches show that 5hmC is not only an intermediate of DNA demethylation, but also plays an important role in many biological processes and human diseases as an epigenetic mediator.^3–6^ The recent development of high-throughput sequencing technology has enabled whole-genome sequencing of 5hmC in mammalian systems. Generally, there are two strategies, which are the selective enrichment-based profiling or deamination based single-base resolution sequencing methods.^7–22^ While the current application of these methods provides key information about the distribution and functional insights of 5hmC, there are major shortage for both strategies. The lack of single-base resolution information of the profiling strategy limited its application in detailed 5hmC location context while the single-base resolution method is limited by its high sequencing cost. Therefore, a method that has both advantage of single-base resolution and enrichment would be highly valued in a broad biological and clinical studies. Recent study developed a bisulfite free strategy to detect 5hmC in single-base resolution using AID/APOBEC family DNA deaminase enzyme as deamination reagent, in which 5hmC is protected by a glucose motif catalyzed by T4 beta-glucosyltransferase (T4-βGT).^23–25^ This strategy has also been reported in a well-studied chemical selective profiling method back to year 2011 using a modified UDP-glucose.^7^

Here we introduce a cost-effective single base resolution 5hmC sequencing method which allows genome-wide chemical labeling and enrichment of 5hmC and a bisulfite free single base resolution detection of 5hmC. Our strategy (DIP-CAB-Seq) has three steps,1) label and enrich the 5hmC containing fragments using the N3-UDP-glucose, click chemistry and biotin-avidin interaction, 2) deaminate unprotected cytosines and 5mC using a deaminase, 3) construct sequencing library in a single-stranded manner. This strategy is compatible with the labeling chemistry, enzyme-based deamination and single-stranded DNA library construction. To demonstrate the superiority and effectiveness of this method, we have applied this approach to compare detected 5hmC signals among our new methods (DIP-CAB-Seq), 5hmC-Seal and ACE-Seq. We found that we can detect reliable single base 5hmC signals using DIP-CAB-Seq method with limited sequencing depth.

## Method

### Cell Culture and DNA preparation

The v6.5 mouse ESCs are the same cell line from a previous study^23^. Cell culture and genomic DNA was prepared as previously described. Briefly, mouse ESCs were cultured on 0.1% gelatin-coated plates in 2i + leukemia inhibitory factor (2i + LIF) media which is consist of N2B27 (DMEM)/F-12, neurobasal, N-2 supplement, and B27 supplement, supplemented with 1 μmol/L PD0325901 (PZ0162, Sigma-Aldrich, St. Louis, USA), 3 μmol/L CHIR99021 (SML1046, Sigma-Aldrich) and 1000 U/mL LIF (PMC9484, Gibco).The genomic DNA was extracted by SDS/proteinase K digestion, phenol/chloroform extraction and ethanol precipitation.

### Library construction

#### Nano-ACE-Seq Library Construction of 5hmC-containing Genomic DNA (DIP-CAB-Seq)

500 ng fragmented mESC genomic DNA (average 200 bp) was treated with β-GT in the presence of UDP-6-N3-Glu and labeled with cyclooctyne-biotin. Subsequently, 5hmC-containing fragments was enriched using Streptavidin Beads. The enriched fragments were subjected to denaturation with NaOH and enzymatic deamination to transform C/mC to U but not hmC. Then the converted DNA was tagged with Illumina compatible adapter and amplified to an appropriate library concentration using Accel-NGS Methyl-Seq DNA Library Kit (Swift Bioscience) combined with NEB Next Multiplex Oligos for Illumina. Libraries were checked for quality and quantified using Agarose Gel Electrophoresis and Qubit3.0 individually.

#### 5hmC-Seal Library Construction of 5hmC-containing Genomic DNA

5hmC-Seal Library was prepared as previously described. Briefly, the fragmented mESC genomic DNA (average 200 bp) was treated with UDP-6-N3-Glu in the presence of β-GT to form chemical modification, followed by label with DBCO-PEG4-Biotin via click reaction. The 5hmC-containing DNA fragments were captured by C1 Streptavidin beads. The beads with enriched DNA fragments were resuspended in water and amplified with 12-17 cycles of PCR using an enzyme mixture in the Nextera kit. The PCR products were purified using AMPure XP beads. Sequencing was performed on the NextSeq instrument.

#### ACE-Seq Library Construction of 5hmC-containing Genomic DNA

ACE-Seq Library was prepared as previously described. Briefly, the m-ESC genomic DNA was sheared with enzymatic to average 200 base pair and purified. DNA fragment was prepared to a concentration of 15~20ng/μL for using. Assemble the reaction in a total volume of 50μL using the table below as a per reaction. If processing multiple samples, making a master mix of everything except the sample should be recommended.

Placed the reaction in thermocycler and incubated at 37□ for 1 hour Subsequent reaction was cleaned separately by DNA Clean & Concentrator-5 Kit (ZYMO Research) and eluted in 25μL EB buffer. Glycosylated dsDNA was treated with a fresh 0.1N NaOH solution, incubated in thermocycler at 50□ for 10 minutes for denaturation. After that, transfer sample into an ice box and stand at least 5 minutes to keep DNA single strand. Followed the ssDNA was subjected to enzymatic deamination to transform C/mC to U but not hmC with NEB Next® Enzymatic Methyl-seq Kit (New England Biolabs). Then the converted DNA was tagged with Illumina compatible adapter and amplified to an appropriate library concentration using Accel-NGS® Methyl-Seq DNA Library Kit (Swift Bioscience) coupled with NEB Next® Multiplex Oligos for Illumina® (New England Biolabs). Libraries were checked for quality and quantified using Agarose Gel Electrophoresis and Qubit3.0 individually.

#### Sequence alignment and peak identification

Sequencing reads were trimmed by trimmomatic with the trim tail option set true, and mapped to the mouse genome (mm10) by bowtie2. The peaks analysis was conducted by MACS2 call peaks, under the pair-end mode with bam file as input format. Afterward, the peaks were annotated by HOMER annotate Peak, and mm10 was used as reference.

#### Single base resolution analysis

The numbers as well as sites information of converted and unconverted cytosines in samples were recognized by bismark, with the criterion that C site cutoff greater than zero, that is, the conversion was valid as long as any read indicates modification of the site.

#### Definition of enhancer subgroups and motif analysis

Enhancers with evidence showing its interaction with distal genomic regions were considered as the most active enhancer subgroup (interacting enhancers). The genomic locations of these enhancers were obtained from published ChIA-PET dataset. The active and poised enhancers were defined by using histone modification markers. Both H3K4me1 and H3K27ac enriched regions were obtained from previous publication. Liftover was used to convert the genomic location from mm8 to mm10. Regions overlapped with interacting enhancers were discarded. Regions enriched with H3K27ac were considered as active enhancers. Regions only enriched with H3K4me1 but not H3K27ac were considered as poised enhancers. To detect TF motifs around 5hmC-modified cytosines, we performed de novo motif analysis with HOMER find Motifs Genome around the 5hmC sites in mouse ESCs.

## Results

### 1. DIP-CAB-Seq can generate deaminated DNA library from the pull-downed DNA fragments

We present here a selective chemical labeling coupled bisulfite free sequencing (DIP-CAB-Seq) approach to generate the genome-wide, single-base resolution maps for 5hmC. In the current approach, we first selectively labeled and enriched 5hmC-containing DNA fragments by using the glucosyltransferase and azido-UDP-glucose based DNA profiling prior to the subsequent deamination (Figure 1). We then performed the deamination on the enriched fragments by using deaminase-based bisulfite free reaction. The normal cytosine and 5mC can be deaminated while 5hmC remained as glucosylated 5hmC (N_3_-5gmC). Therefore, 5hmC will be read as C after PCR amplification while normal C and 5mC will be read as T. After the single stranded library construction and PCR amplification, the 5hmC containing fragments will be selectively amplified while the 5hmC can be read in single-base resolution and generate the precise genomic locations of 5hmC.

**Figure 1.**
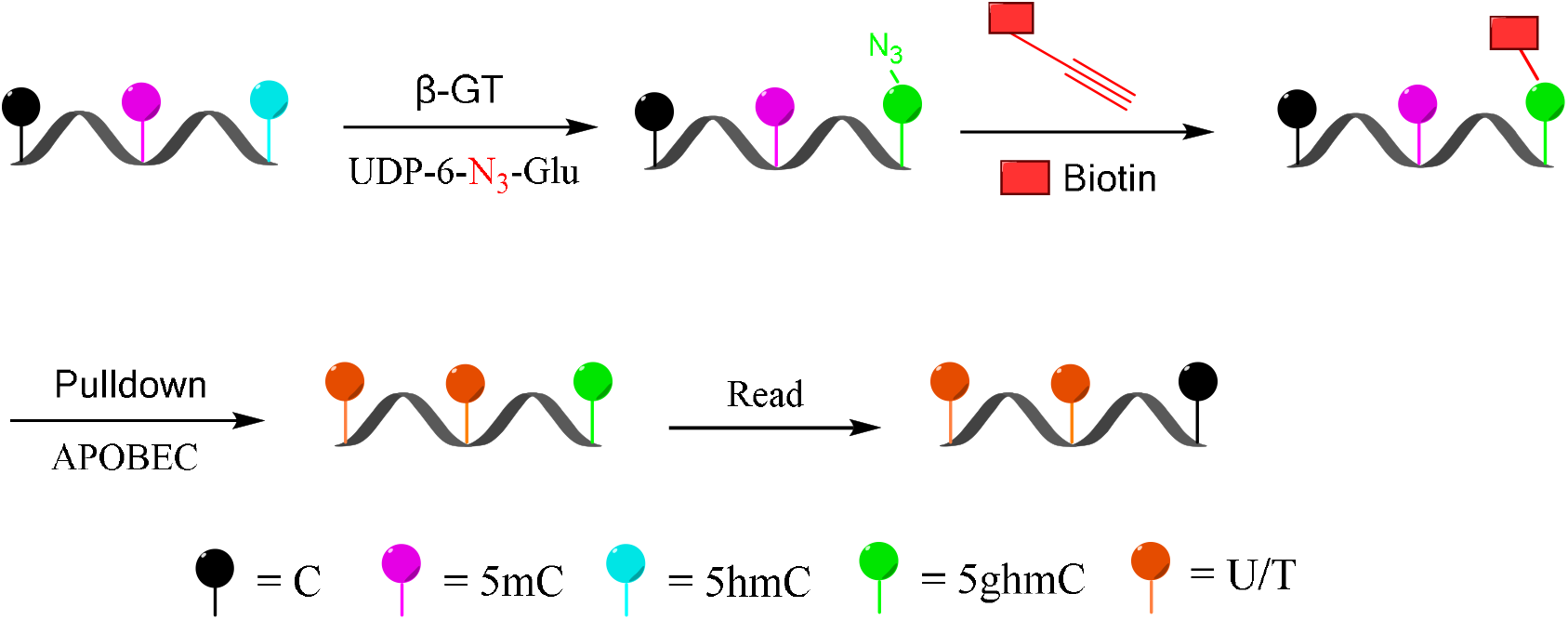
Schematic diagram of selectively labeling and deaminase-based sequencing of 5hmC in DNA (DIP-CAB-Seq).

The first challenge is whether we can generate high quality next-generation sequencing library from the trace amount pull-downed DNA. We validated that approach by generating 5hmC library in genomic DNA of mECS. We constructed ACE-Seq libraries and DIP-CAB-Seq libraries from two replicated mECS gDNA, respectively. The ACE-Seq library generate 35067558 reads with 68% mapping ratio, while the DIP-CAB-Seq library generate 32769029 reads with 61% mapping ratio, not far from the former result. The DIP-CAB-Seq methods have slightly lower mapping ratio than ACE-Seq method which is fully acceptable considering the ultra-low input DNA amount in the library construction step in DIP-CAB-Seq method. Both DIP-CAB-Seq libraries and regular ACE-Seq libraries have similar total sequencing reads which guaranteed us a fair comparison between ability of the two methods in detecting the 5hmC sites in genome. These data indicate that we can successfully construct the 5hmC single-base resolution library from the enriched DNA fragments.

### 2. DIP-CAB-Seq can selectively enrich 5hmC-containing DNA fragments

Using the DIP-CAB-Seq strategy, we performed both regular hmC-Seal profiling and single-base resolution mapping of 5hmC on mESCs genomic DNA. The profiling analysis indicated that 5hmC from both libraries accumulate at intergenic and intron regions in mouse ESCs (Figure 2a). This observation is consistent with previous findings.^7^ We barely observed any abnormal 5hmC signals nor did we observe any noticeable changes of 5hmC at these genomic element regions between the two methods. Therefore, the DIP-CAB-Seq can totally inherit the profiling information that the traditional hmC-Seal method carried.

**Figure 2.**
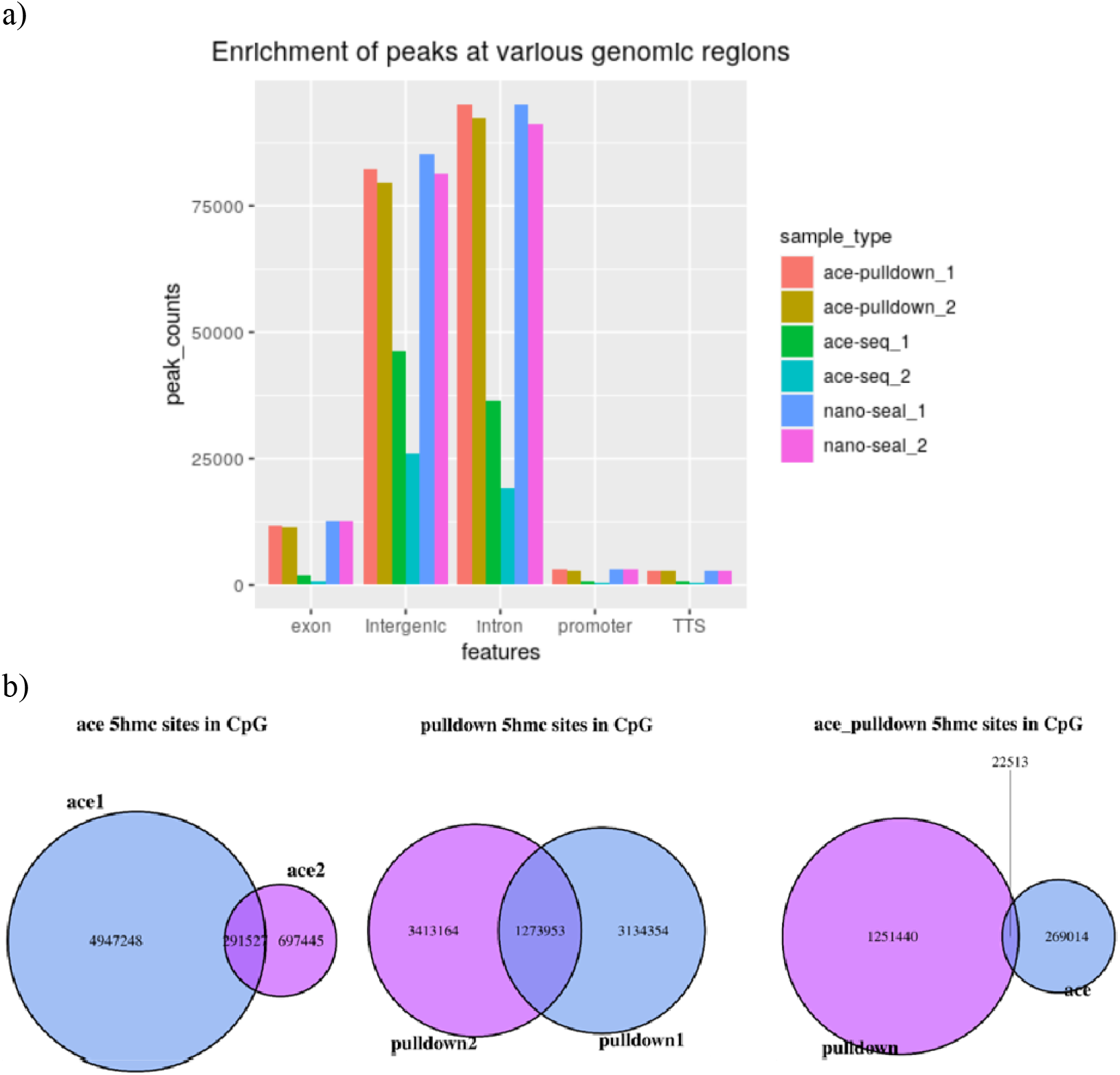
Enrichment of 5hmC-containing DNA fragments with ace, DIP-CAB-Seq and nano-seal methods a) Enrichment of peaks at various genomic regions b) Repeatability of 5hmC sites with ace, DIP-CAB-Seq and nano-seal methods

### 3. DIP-CAB-Seq can detect single base information with limited sequencing depth

We next analyzed 5hmC enriched regions using the base-resolution data in ACE-Seq and DIP-CAB-Seq so as to evaluate the repeatability of both methods. We observed 5936200 5hmC sites in CpG context in the union set of two ACE-Seq samples, while there’re only 291527 5hmC sites, that is, only 4.9% of the union set in the intersection set of both samples. On the other hand, we found 7821453 5hmC sites in CpG context in the union set of the DIP-CAB-Seq samples, and there are 1273953 5hmC sites in the intersection set of both samples, accounting for 16.3% of the union set. These data indicate that the DIP-CAB-Seq method has higher detection ability and consistency than ACE-Seq with same sequencing depth. Considering the total sequencing depth is much lower compared to a regular expensive whole genome 5hmC sequencing, the DIP-CAB-Seq method shows much better application potential in large-scale sample studies. We also detect 5hmC signals in CHH and CHG, which may have 5hmC sites according to the previous study. As expected, DIP-CAB-Seq methods showed higher consistency than ACE-Seq. We observed lots of CHH and CHG sites from the ACE-Seq library, which may come from the false positive signal of the ACE-Seq due to the non-perfect deamination. Therefore, the enrichment of the 5hmC fragments can also decrease the false positive signals even if the deamination reagents is not perfect in a bisulfite free system. We further analyzed the single-base resolution signals from both method in genomic elements.

Based on the above findings, we observed much better consistency from DIP-CAB-Seq method than ACE-Seq, indicating the DIP-CAB-Seq method could have better understanding of the biology with limited sequencing depth (Fig. 2b).

### 4. DIP-CAB-Seq can detect detailed preferential occurrences of 5hmC

With single-base resolution information available, we continued to investigate the detailed preferential occurrences of 5hmC sites in the genome to understand the potential of this method in biology study.

The previous base-resolution mapping of 5hmC allowed for the determination of the surrounding base composition of 5hmC sites in mouse ESCs.^22^ We aligned our 5hmC sites in CG context and examined the base compositions (Fig. S1). Compared to reported results, our data possesses a similar local sequence context, having increased guanine abundance with a depletion of thymine.

We then performed motif analysis with HOMER32 at +/− 100 bp region around the 5hmC sites. We studied the well-known 5hmC containing motif Klf4, Oct4, Hif1a, Esrrb, and Sox2, our motif analysis successfully identified these motifs around the 5hmC sites in mouse ESCs (Fig.3). Our data indicated that our method can obtain these detailed 5hmC signals with very low sequencing depth compared to the reported method.

**Figure 3.**
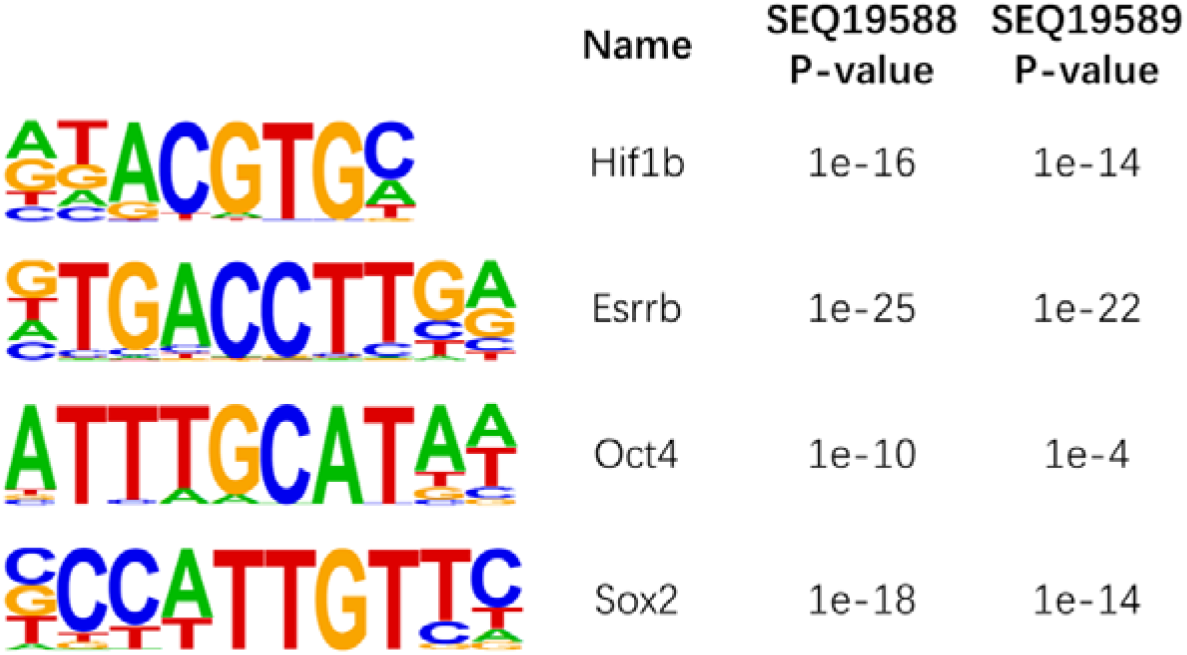
Motif analysis of 5hmC sites in mouse ESCs

## Discussion

The current dilemma between cost and sequencing resolution has hamper lots of biological and clinical studies of 5hmC. Generally, 5hmC has 10 times lower genome percentage than 5mC and 5hmC is distributed broadly across the genome, which requires very deep sequencing for detecting low abundant 5hmC sites. Although the profiling methods provides a cheap manner to map the 5hmC peak, it is much more difficult to quantify the 5hmC signals than the single-base resolution method. DIP-CAB-Seq method we developed can solve this problem, although we cannot quantify the 5hmC abundance for a specific 5hmC site, it provides much better resolution than the regular profiling method. We showed that the base-resolution data from the very limited sequencing depth can provide detailed 5hmC loci information. It is well known that 5hmC profiling has the potential to be a clinical tool in different clinical questions. The current profiling can only calculate the 5hmC score based on 5hmC reads density, which needs lots of adjustment between the samples and cohorts. To further study the role of 5hmC in clinical application, our method can provide better quantification information and could be more reliable than the profiling method.

## Conclusions

Herein, we developed a method based on ESC whole-genome analysis which could enrich 5hmC-containing DNA by selective chemical labeling and locate 5hmC sites at single-base resolution with enzyme-based deamination. The combination experimental design is an extension of previous methods, and we hope that this cost-effective single base resolution 5hmC sequencing method could be used to promote the mechanism and diagnosis research of 5hmC.

## Funding

This research was supported by the Beijing Natural Science Foundation (7194312 to X.L.), Beijing Key Clinical Specialty for Laboratory Medicine-Excellent Project (No. ZK201000) and Beijing Key Laboratory for Mechanisms Research and Precision Diagnosis of Invasive Fungal Diseases.

## Footnote

### Conflicts of Interest

All authors have no conflicts of interest to declare.

### Ethical Statement

None required.

## Contributions

(I) Conception and design: X Li; (II) Administrative support: Y Xu; (III) Provision of study materials: W Guo, X Li; (IV) Collection and assembly of data: X Li, X Shi, Y Gong; (V) Data analysis and interpretation: X Li, X Shi; (VI) Manuscript writing: All authors; (VII) Final approval of manuscript: All authors.

## Notes

### Competing Interest Statement

The authors have declared no competing interest.

### Summary of Updates

Figure 2 revised; author affiliations updated; Supplemental files updated.

