## Supporting Information for "Selective Chemical Labeling and Sequencing of 5-Hydroxymethylcytosine in DNA at Single-Base Resolution"

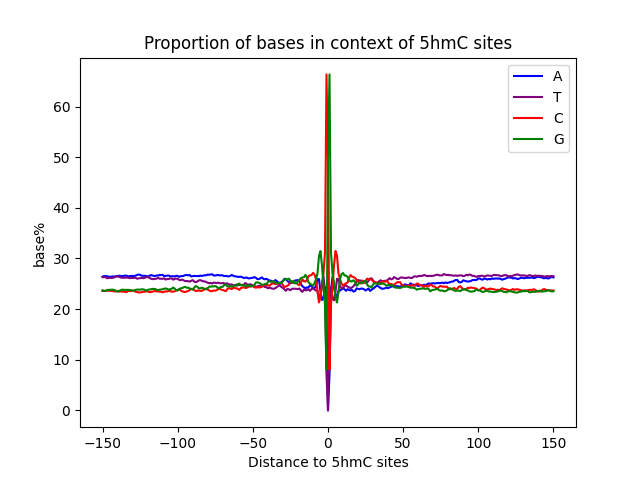

Figure S1 Surrounding base composition of 5hmC sites in mouse ESCs

**Table S1 comparative analysis**

| Method | seq-num | trimmomatic | bam-reads | bowtie2_0times | bowtie2_1times | bowtie2_2times |
| --- | --- | --- | --- | --- | --- | --- |
| nano-seal | SEQ19585 | 0.979 | 32840003 | 0.3021 | 0.5952 | 0.1027 |
| nano-seal | SEQ19590 | 0.9772 | 37038581 | 0.2601 | 0.6298 | 0.11 |
| ace-seq | SEQ19586 | 0.9876 | 34104219 | 0.2529 | 0.6746 | 0.0725 |
| ace-seq | SEQ19587 | 0.9875 | 36030896 | 0.2455 | 0.6839 | 0.0706 |
| ace-pulldown | SEQ19588 | 0.8946 | 34705347 | 0.3574 | 0.5883 | 0.0543 |
| ace-pulldown | SEQ19589 | 0.9198 | 30832711 | 0.3114 | 0.6294 | 0.0592 |

Table S2 Statistics results of 5hmC sites

| Method | seq_num | total_lines | total_strings | total_C | hmC_CpG | hmC_CHG | hmC_CHH | C2T_CpG | C2T_CHG | C2T_CHH |
| --- | --- | --- | --- | --- | --- | --- | --- | --- | --- | --- |
| ace-seq | SEQ19586 | 18191578 | 36383156 | 756963288 | 6342475 | 20163184 | 62652700 | 27525310 | 142500601 | 497779018 |
| ace-seq | SEQ19587 | 18824449 | 37648898 | 812058698 | 1183130 | 3235097 | 11574624 | 32824248 | 170305710 | 592935889 |
| ace-  pulldown | SEQ19588 | 12680153 | 25360306 | 614206950 | 5427793 | 1418759 | 4081655 | 41829555 | 144165807 | 417283381 |
| ace-  pulldown | SEQ19589 | 12522156 | 25044312 | 598297204 | 5818168 | 2586082 | 7293592 | 41661802 | 140467042 | 400470518 |
